# Three Distinct Annotation Platforms Differ in Detection of Antimicrobial Resistance Genes in Long-Read, Short-Read, and Hybrid Sequences Derived from Total Genomic DNA or from Purified Plasmid DNA

**DOI:** 10.1101/2022.07.27.501810

**Authors:** Grazieli Maboni, Rodrigo de Paula Baptista, Joy Wireman, Isaac Framst, Anne O. Summers, Susan Sanchez

## Abstract

Recent advances and lower costs in rapid high-throughput sequencing have engendered hope that whole genome sequencing (WGS) might afford complete resistome characterization in clinical bacterial isolates. Despite its potential, several challenges should be addressed before adopting WGS to detect antimicrobial resistance (AMR) genes in the clinical laboratory. Here, with three distinct ESKAPE bacteria, we compared different approaches to identify best practices for detection of AMR genes, including: total genomic DNA and plasmid DNA extractions, solo assembly of Illumina short-reads and of ONT long-reads, two hybrid assembly pipelines, and three *in silico* AMR databases. We also determined the susceptibility of each strain to 21 antimicrobials. We found that all AMR genes detected in pure plasmid DNA were also detectable in total genomic DNA indicating that, at least in these three enterobacterial genera, purification of plasmid DNA was not necessary to detect plasmid-borne AMR genes. We also found that Illumina short-reads used with ONT long-reads in either hybrid or polished assemblies of total genomic DNA enhanced sensitivity and accuracy of AMR gene detection. Phenotypic susceptibility corresponded well with genotypes identified by sequencing, but the three AMR databases differed significantly in distinguishing mobile dedicated AMR genes from non-mobile chromosomal housekeeping genes in which rare spontaneous resistance mutations might occur. This study reveals the need for standardized biochemical and informatic procedures and database resources for consistent, reliable AMR genotyping to take full advantage of WGS to expedite patient treatment and to track AMR genes within the hospital and community.

## 1. INTRODUCTION

Antimicrobial resistance is increasingly threatening global public health. Current routine drug susceptibility testing uses cultivation-based phenotyping with several commercial automated platforms [1]. Nonetheless, phenotyping may take three to four days for fast-growing and weeks for slow-growing bacteria [2, 3] and considerably longer in non-automated laboratories. Interpretation of susceptibility data is challenging since the lack of clinically relevant breakpoints for several pathogens [2-4] can delay decisions on adequate antimicrobial therapy.

Recent technical improvements and lower costs in rapid high-throughput DNA sequencing have engendered hope that whole genome sequencing (WGS) might afford rapid detection of clinically relevant resistance genes. Consequent accurate prediction of resistance phenotypes could complement or even replace slower cultivation-based tests [5]. Genomic data has correctly predicted phenotypic resistance for some antimicrobial resistance (AMR) genes with established susceptibility breakpoints based on minimum inhibitory concentration (MIC) of the corresponding antimicrobial [5-10]. However, WGS protocols for accurately detecting AMR genes are not yet standardized. This multistep process includes specimen propagation in suitable medium, DNA extraction, sequencing library preparation, generation of sequence ‘reads’, assembly of reads into chromosome or plasmid sequences, and identification of antimicrobial resistance genes [11]. Since clinically relevant AMR genes are often carried on mobile genetic elements (transposons, integrons and plasmids) effective detection of these widely transmissible elements is essential. Purification of plasmid DNA is not as simple as that of total cellular (aka genomic) DNA, and there was concern that the latter might inefficiently recover large, low copy number plasmids [11-13], impairing detection of plasmid-borne AMR genes.

Sequence acquisition platforms and assembly pipelines deeply influence the fidelity of the final assembly and thus the accurate annotation of all genes encoded by chromosomes and plasmids [5]. The current industry standard approach to full genome sequencing combines the highly accurate Illumina short-read sequencing with the newer long-read platforms such as Oxford Nanopore Technologies (ONT) or PacBio which can better distinguish separate instances of repeated loci, the *bete noire* of genome assembly, especially in prokaryotes [14]. Nanopore’s long-read capacity is a real boon to sequencing plasmids, whose frequent repeated regions thwart correct computational assembly of short-read data delivered by Illumina [9]. This advantage also applies to the assembly of bacterial whole genomes [14] because long-reads enable the correct structural resolution of complex genomic regions. The main drawback of using ONT sequencing alone is the error rate of raw sequence reads when compared to the more precise Illumina short-read technology [15]. Thus, combining short- and long-read sequencing has become best practice for sequencing typically closed circular prokaryotic cell chromosomes (∼4-5 Mb in enterobacteriaceae) and plasmids (ranging from 2-800 kb, also typically double stranded closed circles) [14, 16, 17] to optimize accurate annotation of all encoded genes including AMR genes.

Finally, identification of AMR genes requires reliable annotation databases of previously sequenced strains including laboratory phenotypic data on their antimicrobial susceptibility. There are three frequently cited curated public databases dedicated only to AMR genes, each using different informatic strategies and data sources [5], Comprehensive Antibiotic Resistance Database (CARD) [18], ResFinder [19], and AMRFinder [20]. We found the outputs from these AMR databases often disagree with each other and with laboratory-based phenotyping. This final all-important step is also very much in need of standardization.

To devise a WGS best practice for timely and accurate detection of relevant AMR genes in clinical isolates we investigated whether such genes were better detected (i) in whole genome DNA or in purified plasmid DNA and (ii) by Nanopore long-read or Illumina short-read or a combination (hybrid). Then, for a given genome or plasmid sequence we asked (iii) which public AMR reporting platform best identified clinically and epidemiologically relevant AMR genes and (iv) did the genotype reported by an AMR database platform correlate with the laboratory-determined susceptibility phenotype of each strain.

## 2. MATERIALS AND METHODS

### 2.1 Bacterial strains

Two clinical bacterial strains, *Enterobacter ludwigii* (LST1391B) and *Klebsiella pneumoniae* (LST1504-C2) were isolated in the late 1970’s at University of Georgia from MacConkey agar plates inoculated with fecal swabs from two unrelated hospital patients provided by the Stuart Levy lab at Tufts University Medical Center in Boston, MA, and were cryopreserved since then. As a control we used the cryopreserved standard *Escherichia coli* laboratory strain (DU1040) carrying the extensively characterized 94-kb conjugative IncFII plasmid, NR1 [21]. The cryopreserved strains were revived by streaking on 5% sheep blood agar (Remel, San Diego, CA, USA) and incubated for 24 hours at 35° C with 5% CO2.

### 2.2 Antimicrobial susceptibility testing

The minimum inhibitory concentration (MIC) of antimicrobials was determined using two systems, Vitek-2 (bioMérieux, Marcy l’Etoile, France) and Trek Sensititre (Trek Diagnostic Systems, Cleveland, OH, USA). For the Vitek-2 testing, three different MIC cards (GN-98, GN-69, and GN-82) were run according to the manufacturer’s instructions (Vitek-2, bioMérieux). For the Trek Sensititre testing, both GN4F and COMPGN1F Gram-negative microplates were run according to the manufacturer’s instructions (Trek Diagnostic Systems). Twenty-one antimicrobials representing eight chemical classes of drugs were tested: lactams and lactamase inhibitors (ampicillin, amoxicillin/clavulanic acid, piperacillin/tazobactam, cefalexin, ceftriaxone, cefazolin, cefepime, ceftazidime); fluoroquinolones (ciprofloxacin, levofloxacin, enrofloxacin); aminoglycosides (gentamicin, amikacin); tetracyclines (doxycycline, tetracycline); antifolates (trimethoprim/sulfamethoxazole), carbapenems (ertapenem, imipenem, meropenem), phenicol (chloramphenicol), and nitrofurans (nitrofurantoin). The resulting MIC value was assigned to clinical categories of susceptible or resistant according to the Clinical & Laboratory Standards Institute (CLSI M-100) [2] and the European Committee on Antimicrobial Susceptibility Testing (EUCAST) [3]. MIC results in “intermediate” or “resistant” ranges were both assigned as “resistant”.

### 2.3 Extraction of total genomic DNA and pure plasmid DNA

To expedite the plasmid purification, we used fresh colonies from non-selective agar rather than a liquid broth culture. Specifically, cryopreserved bacterial cells were streaked for colony isolation on Luria-Bertani agar without antibiotics (Remel) and incubated for 18 hours at 35° C with 5% CO2. All growth on agar was scraped from the plate surface, transferred into 500 mL of Luria-Bertani broth (Remel), and incubated at 35° C, at 120 rpm for 3 to 4 hours until approximately mid-exponential phase (OD_600nm_ = 0.6). Then the entire culture was centrifuged at 6000 rpm for 15 minutes at 4°C; the supernatant spent medium was discarded, and plasmids were extracted from the cell pellet with the Qiagen Large Construct Kit. According to the manufacturer’s instructions, total DNA extraction was performed on this suspension of freshly grown bacterial colonies using the Genomic-tip 500/G kit (Qiagen, Hilden, Germany). Plasmids were electrophoresed in a 0.5% SeaKem Gold agarose with a Tris-acetate buffer, pH 8 (Lonza), stained with SyberGreen (Sigma Aldrich), and imaged as previously described [21]. Plasmid molecular weight was estimated by a semi-log polynomial fit of the migration distances of standard supercoiled plasmid DNA bands of known molecular mass.

The concentration of total genomic or plasmid DNA preparations was quantified by a Qubit 2.0 fluorometer using a double-stranded DNA assay kit. Purity was assessed by NanoDrop Spectrophotometer, according to the manufacturer’s instructions (Thermo Scientific, Waltham, MA, USA). DNA preparations were stored at -20 °C until sequencing.

### 2.4 Whole-genome and plasmid sequencing

MinION libraries were prepared from 400 ng of pure plasmid or total genomic DNA using the SQK-RBK004 Rapid Barcoding Kit and sequenced with the FLOW-MIN 106 (R9.4 SpotOn) flow cell, according to instructions from ONT. Total genomic DNA and pure plasmid DNA of each strain were barcoded separately. ONT’s MinKNOW software (version v18.03.1) collected raw electronic data as Fast5 read files and bases were called by ONT’s EPI2ME software. Initial real-time workflows “What Is in My Pot?” (WIMP) were used to confirm bacterial biotype based on 16S rDNA. Sequence data were collected for 24 hours.

Illumina paired-end libraries were prepared from total genomic DNA or pure plasmid DNA using a Nextera DNA Flex library prep kit on an Illumina iSeq 100 instrument and sequenced with 150 bp paired reads according to the manufacturer’s instructions. Quality control of library preparation was performed using the QIAxcel Advanced Systems (Qiagen, Hilden, Germany). Nanopore and Illumina sequencing was performed at the Athens Veterinary Diagnostic Laboratory, University of Georgia, USA.

### 2.5 Contig assembly and data analysis

All raw reads were submitted to a metagenomics pipeline to detect species or potential contamination using Metaphlan2 v2.7.8 before the *de novo* assembly [22]. Each genome or plasmid was assembled using four different strategies: (i) Nanopore long-reads, named in this study as Nanopore-only approach; (ii) Illumina short-reads, named in this study as Illumina-only approach; (iii) simultaneous assembly of Illumina reads and Nanopore reads, named as Hybrid approach; (iv) and Flye-assembled Nanopore reads post-hoc matched with Illumina reads, named as Nanopore-polished approach. Assembly was guided by the average published chromosome size for the respective bacterial genus available in GenBank. The assembly of pure plasmid sequences was guided by the estimated molecular size of the plasmids observed in agarose gel electrophoresis (**Figure 1**).

**Figure 1.**
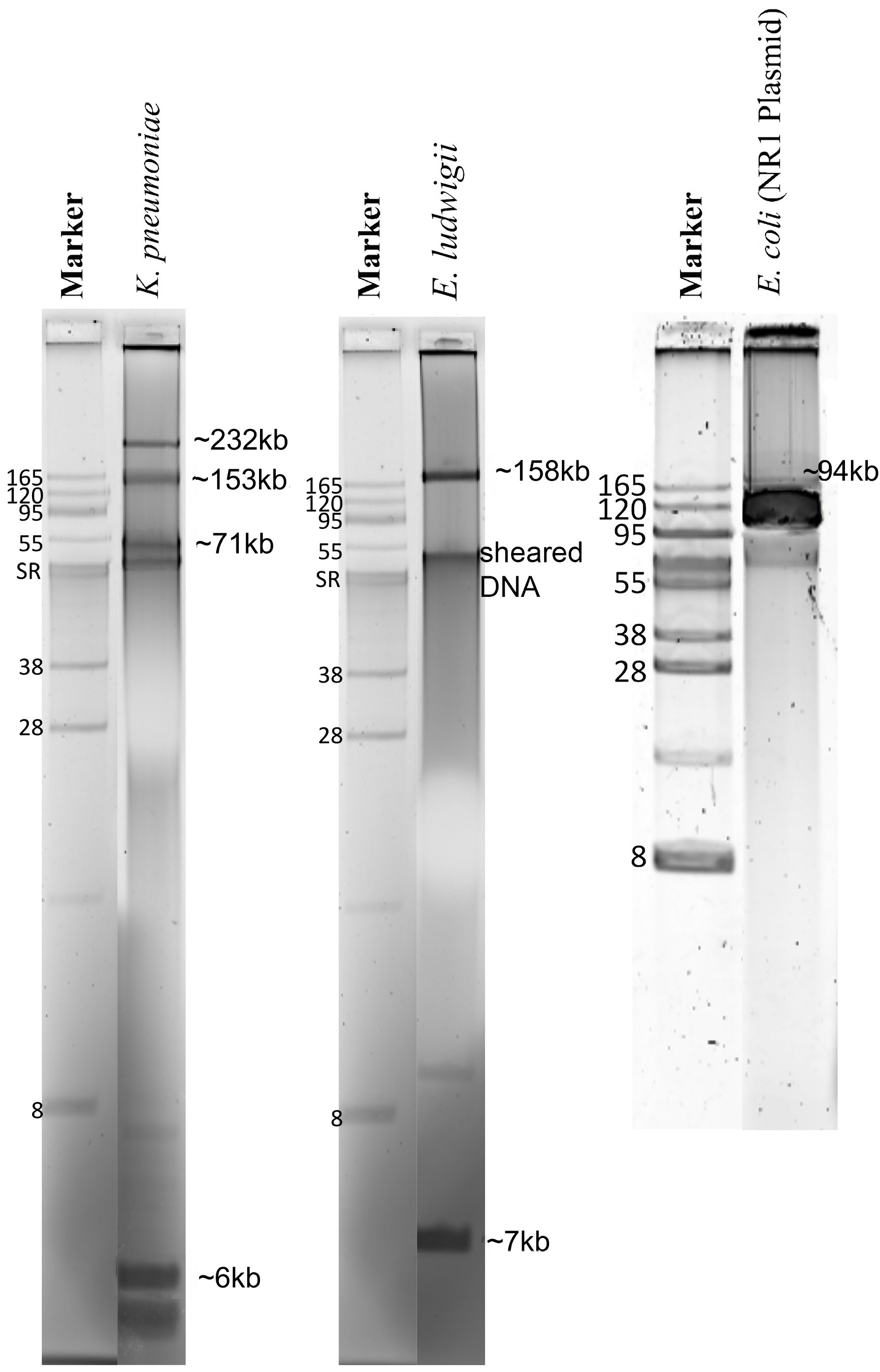
Agarose electrophoresis of plasmids from three enterobacteria sequenced in this study. The gel is 0.5% SeaKem Gold agarose stained with SyberGreen. Blue boxes highlight the plasmid DNA bands. Plasmid molecular weight (kilobases= kb) was estimated by semi-log plotting based on DNA band sizes observed in the agarose gel electrophoresis. Strains: *Klebsiella pneumoniae* - LST1504-C2, *Enterobacter ludwigii* -LST1391B, and *Escherichia coli* - DU1040 (NR1) plasmid control.

For the Nanopore-only assembly, barcoded sequencing reads were demultiplexed using Porechop v0.2.4 (https://github.com/rrwick/Porechop) and assembled using Flye v2.6 [23]. Potential base-call errors in the assembly were verified by Racon v.1.4.7 [24], which generates a genomic consensus with better quality than the output generated by assembly methods using the alignment coverage of the contig blocks. An additional correction step was made in the previous Nanopore-only assembly using Illumina reads for the Nanopore-Illumina polished approach. This correction was done by combining two alignment iterations using BWA v0.7.17 [25] with the MEM function and SMALT v0.7.4 (https://www.sanger.ac.uk/tool/smalt-0/); the output alignment was submitted to the polishing tool Pilon v.1.23 [26].

Besides Flye v.2.6, assembly of plasmid raw reads generated by ONT was also performed using Canu v.2.2 [27]. For Nanopore-only plasmid assembly with Canu, trimmed reads from Porechop were input to Canu v.2.2 using the default options for ONT reads. For Nanopore-polished plasmid assemblies, BWA-MEM v0.7.17 [25] was used to generate alignment overlaps between the ONT plasmid draft assembly and the Illumina reads generated from the plasmid prep. The alignment was than parsed using Samtools v1.12 [28] and used for base call polishing in two rounds of Pilon v1.23 [26].

All Illumina paired end read sequences generated were quality checked with FastQC and trimmed by Trimomatic v.0.36 [25] to remove sequencing adapters and reads with Phred scores < 30. The *de novo* assembly of Illumina reads was performed using Spades v3.12 [29] and polished by Pilon v.1.23 using the same Illumina alignment protocol applied for the Nanopore-Illumina polished approach described above. For the Illumina-hybrid approach, both Illumina and Nanopore reads were submitted to a *de novo* assembly using hybridSPAdes [30] incorporated in Spades v3.12 and were polished by Pilon. This approach prioritizes the contig formation by using de Bruijn graphs with the Illumina short-reads and then mapping long-reads in the edges of the assembly graph to increase contiguity and generate longer scaffolds. PlasmidSPADES, also available in Spades v3.12, was used to optimize plasmid contig assembly from the Illumina data [31]. Taxonomic classification of plasmids was performed using COPLA v1.0 [32]. Assembly statistics were assessed by QUAST v5.0.2 [33]. Annotations of chromosomes and plasmids were performed using Prokka v1.13 [34].

Our bioinformatic pipeline shell script with dependencies, install instructions and usage instructions is available at https://github.com/iframst/HybridAMRgenotyping

### 2.6 Identification of AMR genes in whole genome or pure plasmid sequences

AMR genes were identified in the whole genome and plasmid sequences using three databases (i) ResFinder for acquired AMR genes, with default settings of 90% nucleotide similarity and a 60% minimum length [35]; (ii) Comprehensive Antibiotic Resistance Database (CARD) with criteria selected as: perfect and strict hits only, excluded nudging of ≥ 95% identity loose hits to strict, and high quality/coverage sequences [18], (iii) AMRFinder from the National Center for Biotechnology Information (NCBI) with minimum BLAST identity cut-off of >90% and >50% alignment coverage; and organism search for optimal analysis of *E. coli* and *K. pneumoniae*. Venn diagrams were performed using Venny (version 2.1.0) [36].

## 3. RESULTS

### 3.1 Concentration and purity of total genomic DNA and pure plasmid DNA

The Genomic-tip 500/G kit protocol yield total genomic DNA with concentrations and purity as described in **Table S1**. We adapted a commercial kit for plasmid DNA extraction from liquid media to purify plasmids from bacterial colonies on an agar plate, eliminating 18 hour overnight growth in liquid medium yielding microgram amounts of pure plasmid DNA in 6 hour (**Table S1; Figure 1**) as demonstrated by our positive control strain with the well-characterized 94 kb *E. coli* plasmid NR1 [21]. By gel electrophoresis these plasmid DNA preparations showed the *E. ludwigii* had two plasmids of ∼158.6 kb and ∼7 kb and the *K. pneumoniae* had four plasmids of ∼232 kb, ∼153 kb, ∼71 kb, and ∼6 kb (**Figure 1**).

### 3.2 Assembly and assessment of total genomic DNA preparations

Our analytical pipeline (**Figure 2**) reports total reads, the average read length, and the total base pairs detected (**Table S2)**. QUAST (**Table S3)** reported chromosomes of the expected size for each strain by four distinct assembly approaches. Expected differences for long and short-read chemistries were high N50’s, low L50’s and low N’s indicating long-read Nanopore assemblies were less fragmented and more contiguous than the short-read Illumina assemblies. Total genomic coding sequences (CDS) annotated by Prokka in Nanopore-only assemblies **(Table S4)** exceeded those of the GenBank reference sequences, but as expected Nanopore-polished, Illumina-only, and Illumina-hybrid genomes agreed better with reference sequences (**Table S4**). The Nanopore-only CDS discrepancy likely reflects frameshifts known to occur in ONT base-calling that were corrected by polishing with Illumina reads.

**Figure 2.**
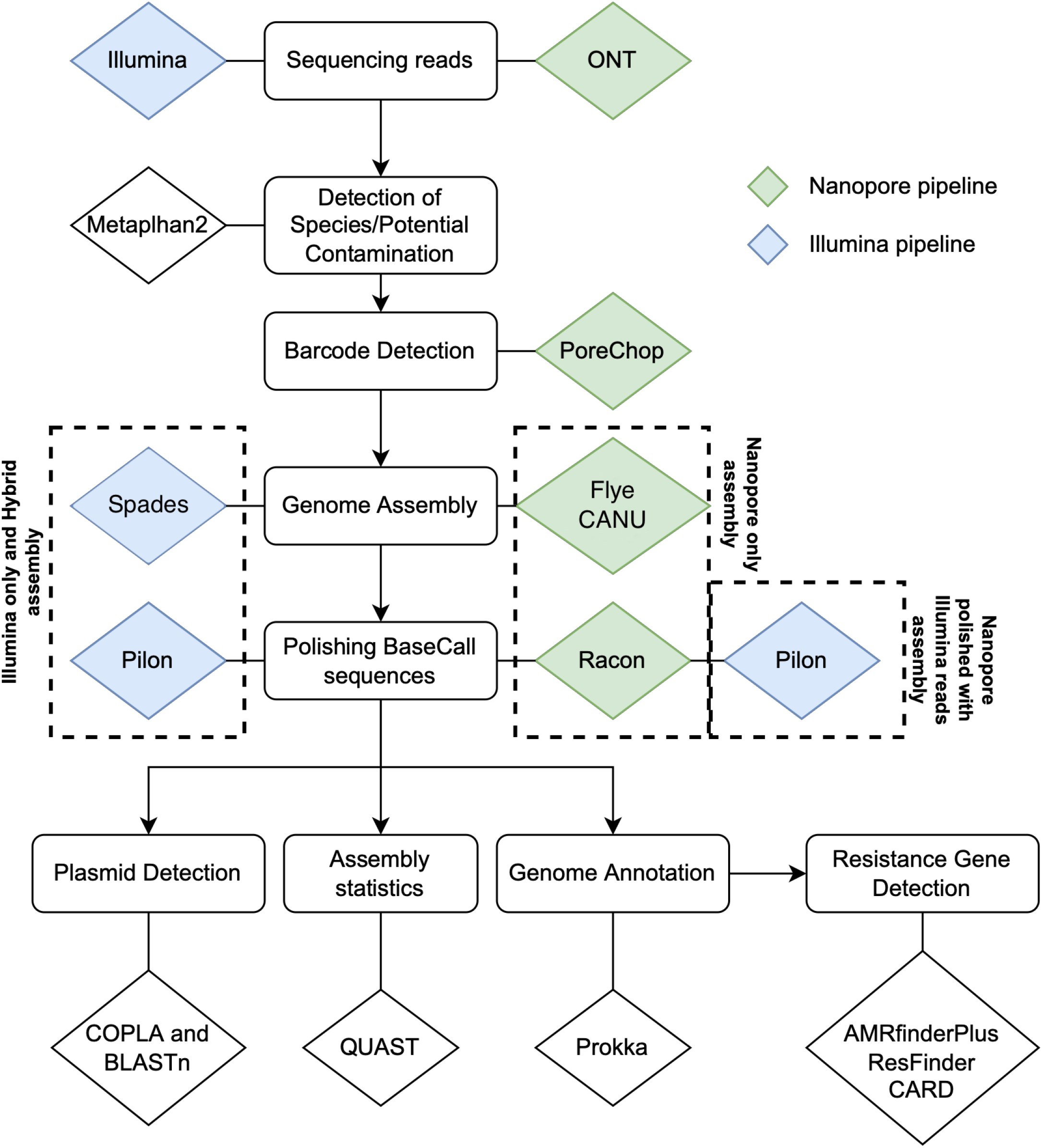
The bioinformatics pipeline used for total genomic and plasmid DNA sequencing analysis. Total genomic DNA and plasmid-only DNA were sequenced using two sequencing chemistries (Illumina and Oxford Nanopore Technologies (ONT)). Reads were assembled using four different approaches: Nanopore-only: MinION reads assembled with Flye, without error correction by Illumina reads; Nanopore-polished: MinION reads assembled with Flye, polished with Illumina reads; Illumina-only: Illumina reads assembled with SPADES, without error correction with Nanopore reads. Illumina-hybrid: assembly of Illumina reads and Nanopore reads with hybridSPADES, polished with Illumina reads. Blue triangles: Illumina. Green triangles: ONT. AMR genes were identified by three different databases: AMRfinder database [20], Resfinder database [35], and CARD [18]. Metaphlan2 [22], Porechop (https://github.com/rrwick/Porechop), Flye [23], Spades [29], Canu [27], Pilon [26], Racon [24], PlasmidSPAdes [40], Prokka [34], Copla [41].

### 3.3 Plasmid sequences assembled from purified plasmid DNA versus total genomic DNA

For the *E. coli* (NR1) pure plasmid DNA sequences, Illumina-hybrid assembly generated a single closed contig of 94,308 bp corresponding to the previously determined 94 kb NR1 plasmid [21]. In contrast, two or several linear contigs of different sizes resulted from the Nanopore-only, Illumina-only, and Nanopore-polished assemblies (**Table 1**). For the *E. ludwigii* pure plasmid DNA sequences, all four assembly approaches generated two linear contigs of ∼130 kb and ∼5 kb consistent with the ∼158.6 kb and ∼7 kb closed supercoiled plasmids seen by electrophoresis. For *K. pneumoniae* pure plasmid DNA sequences, four linear contigs were obtained with Nanopore-only and with Nanopore-polished assemblies, roughly corresponding to the supercoiled bands consistent with the ∼232 kb, ∼153 kb, ∼71 kb, and ∼6 kb visible in the gel. The smallest plasmid of *K. pneumoniae* assembled into a single linear contig of 3-5 kb by all assembly approaches roughly similar to the ∼6 kb supercoiled plasmid in the electrophoresis gel (**Table 1**).

**Table 1.** Plasmid sequences inferred from Nanopore and/or Illumina sequencing assemblies of pure plasmid DNA or of total genomic DNA.

| Strain | Known or<br>Estimated<br>Plasmid Size (kb) | Nanopore-only<sup>a</sup><br>Contig (bp) | Nanopore-polished<sup>b</sup><br>Contig (bp) | Illumina-only<sup>c</sup><br>Contig (bp) | Illumina-hybrid<sup>d</sup><br>Contig (bp) |
| --- | --- | --- | --- | --- | --- |
| **Pure Plasmid DNA** |  |  |  |  |  |
| *E. coli* DU1040 (NR1) | 94.289 | Several contigs<sup>e</sup> | 93,868 | Several contigs | 94,308 |
| *K. pneumoniae*<br>LST1504-C2 | 232 | 313,736 | 313,014 | Several contigs | Several contigs |
|  | 153 | 265,010 | 265,397 | Several contigs | Several contigs |
|  | 71 | 58,149 | 58,214 | Several contigs | Several contigs |
|  | 6 | 3,380 | 3,378 | 3,003 | 6,261 |
| *E. ludwigii* LST1391B | 158.6 | 133,397 | 133,378 | 130,070 | 130,070 |
|  | 7 | 5,187 | 5,152 | 5,283 | 5,152 |
| **Total Genomic DNA** |  |  |  |  |  |
| *E. coli* DU1040 (NR1) | 94.289 | 94,296 | 94,410 | Several contigs | 93,656 |
| *K. pneumoniae*<br>LST1504-C2 | 232 | 313,645 | 312,971 | Several contigs | 314,384 |
|  | 153 | 264,823 | 264,679 | Several contigs | Several contigs |
|  | 71 | 58,157 | 58,118 | Several contigs | Several contigs |
|  | 6 | 3,350 | 3,326 | 2,930 | 2,930 |
| *E. ludwigii* LST1391B | 158.6 | 133,990 | 133,333 | 130,070 | 130,070 |
|  | 7 | 6,174 | 6,174 | Several contigs | 5,033 |

<sup>a</sup>Nanopore-only: MinION reads assembled with Flye, without error correction by Illumina reads.
<sup>b</sup>Nanopore-polished: MinION reads assembled with Flye, polished with Illumina reads.
<sup>c</sup>Illumina-only: Illumina reads assembled with SPADES, without error correction with Nanopore reads.
<sup>d</sup>Illumina-hybrid: assembly of Illumina reads and Nanopore reads with hybridSPADES, polished with Illumina reads.
<sup>e</sup>Several contigs: two or more contigs of different lengths matched with the control or with a plasmid sequence from GenBank.

The total genomic DNA preparations afforded assembly of individual linear plasmids via the Nanopore-only and the Nanopore-Illumina and hybrid methods for all bacterial strains (**Table 1**). As expected, Nanopore long-reads assembled into single linear contigs with sizes corresponding approximately to those observed in the purified plasmid preparations and on the corresponding gels, whereas Illumina short-reads assembled into several linear contigs of miscellaneous sizes.

The COPLA Taxonomic Classifier for plasmids [32] correctly identified the NR1 control as incompatibility group (IncFII), mobility class (MOBF), and mating pair formation (type F) (**Table S5**) placing it in the Plasmid Taxonomic Unit, PTU-FIIE. Taxonomic classifications of the two *E. ludwigii* and four *K. pneumoniae* plasmids were revealed by COPLA to belong to PTU-E3, and PTU-HI1B and PTU-E71III, respectively (**Table S5)**.

### 3.4 AMR genes detected in total genomic DNA *versus* plasmid DNA

We further evaluated whether a plasmid extraction step must be incorporated in a sequencing workflow to obtain reliable identification of AMR genes. For that, we compared AMR genes obtained from plasmid extraction only against those from total genomic extractions. All AMR genes detected in the plasmid extractions were also present in the total genomic DNA extractions of all bacterial strains investigated (**Table 2**). Further, more AMR genes were detected in total genomic than in plasmids-only independent of the assembly approach applied (see **Tables S6-8** for a complete list of genes). In *K. pneumoniae*, genes expected to be detected in plasmids were only found in total genomic DNA (i.e., *aadA1* and *qacEdelta1*), which could be due to unidentified issues in the extraction, sequencing chemistry, or assembling (**Table 2**). Our findings indicate that plasmids were extracted and sequenced along with chromosomal DNA using this study’s total genomic DNA protocol. As plasmid extraction, sequencing, and assembling protocols were cumbersome and did not provide any additional information from what we found in total genomic DNA extraction, we suggest that a plasmid DNA extraction/sequencing step may not be essential to obtain a complete list of AMR genes.

**Table 2.** Antimicrobial resistance (AMR) genes detected in the total genomic *versus* in pure plasmid DNA preparations^*a*^.

| ResFinder |  | AMRFinder |  |
| --- | --- | --- | --- |
| Total Genomic | Pure Plasmid | Total Genomic | Pure Plasmid |
| *Escherichia coli* DU1040 (NR1) |  |  |  |
| *aadA1* | *aadA1* | *aadA1* | *aadA1* |
| *catA1* | *catA1* | *catA1* | *catA1* |
| *sulI* | *sulI* | *sulI* | *sulI* |
| *tet(B)* | *tet(B)* | *tet(B)* | *tet(B)* |
| *mdfA* |  | *qacEdelta1* | *qacEdelta1* |
|  |  | *blaEC* |  |
| *Enterobacter ludwigii* LST1391B |  |  |  |
| *blaACT-12* | *blaACT-12* | *blaACT* | *blaACT* |
| *fosA2* | *fosA2* | *fosA2* | *fosA2* |
|  |  | *oqxA* | *oqxA* |
|  |  | *oqxB* | *oqxB* |
| *Klebsiella pneumoniae* LST1504-C2 |  |  |  |
| *blaSHV-40* | Not identified<sup>b</sup> | *blaSHV* | Not identified<sup>b</sup> |
| *fosA* |  | *fosA* |  |
|  |  | *aadA1* |  |
|  |  | *qacEdelta1* |  |

<sup>a</sup> Results based on Illumina-hybrid assemblies which exhibited more AMR genes. See Methods and Table 1 footnote for details of assembly pipelines.
<sup>b</sup> Not identified: no antimicrobial resistance genes were detected.
**Bold:** genes detected in plasmids observed in both the pure plasmid and total genomic DNA preparations.
Aminoglycoside: *aadA1*. Beta-lactams: *blaACT*, *blaSHV*, *blaEC*. Phenicol: *catA1*. Fosfomycin: *fosA2*. Macrolide, tetracycline, phenicol (broad spectrum): *mdfA*. Tetracycline: *tet(B)*. Sulphonamide: *sulI*. Phenicol/Quinolone: *oqxA*, *oqxB*. Quaternary ammonium: *qacEdelta1*.

### 3.5 Phenotypic *versus* genotypic antimicrobial susceptibility

As discrepancies observed between phenotypic results and the expected genomic outcome are often caused by incorrect susceptibility testing [5], we used two different methodologies to ensure the predicted phenotypes. MIC interpretation of Vitek-2 and Sensititre systems were consistent, but a few discrepant results were observed in *E. ludwigii* and *K. pneumoniae*. For instance, *E. cloacae* MIC to chloramphenicol indicated intermediate resistance by VITEK-2, while by Sensititre it was susceptible. Similarly, MIC of *K. pneumoniae* to amoxicillin/clavulanic acid was susceptible by Vitek-2 but resistant by the Sensititre (**Table 3**).

**Table 3.**
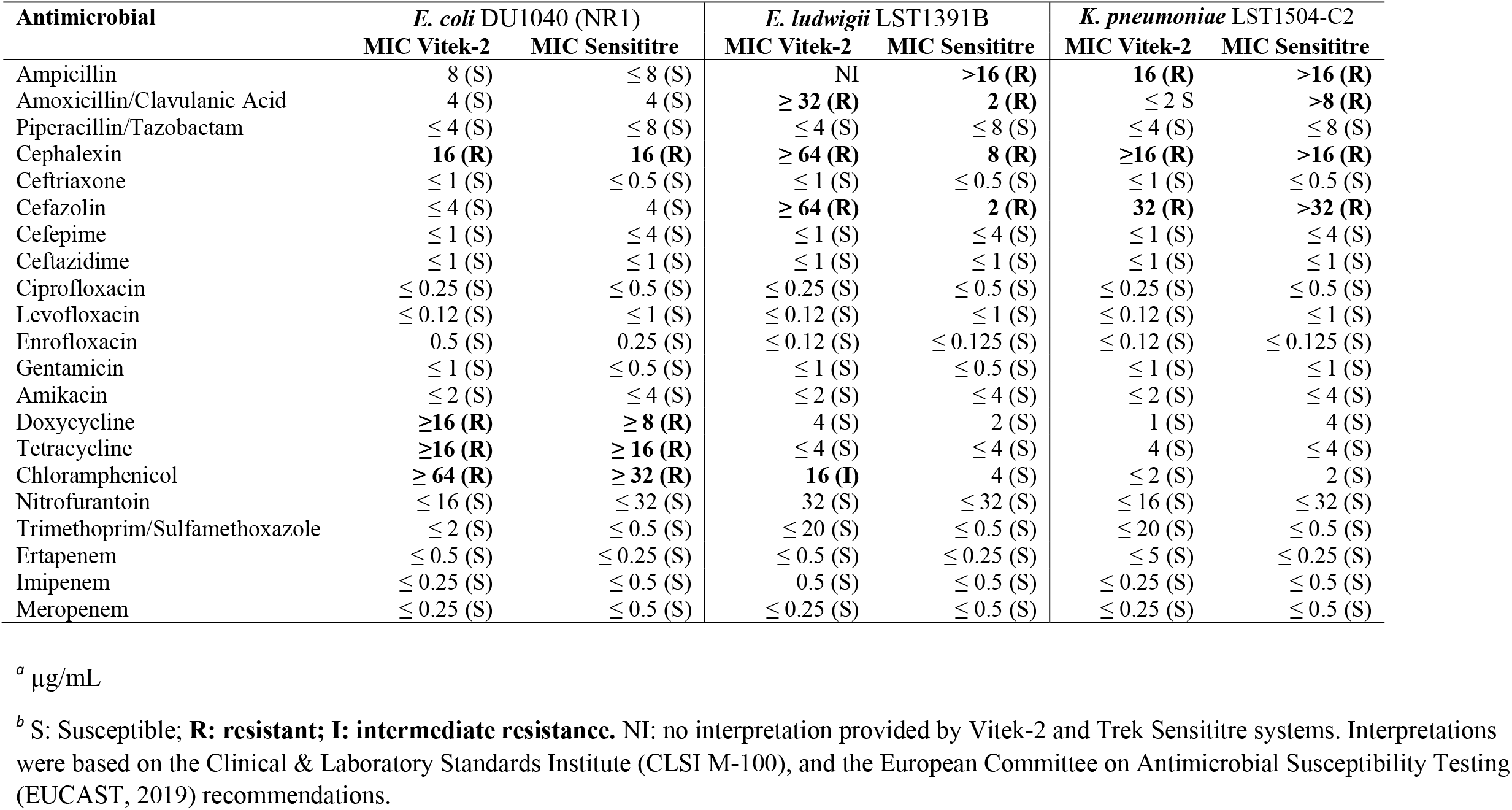
Minimum inhibitory concentration (MIC) ^*a*^ of antimicrobials for the standard and two test strains by two methods ^*b*^.

We then evaluated the ability of the WGS data to identify AMR genes associated with a resistant phenotype correctly. For that, phenotypic resistance was compared with the predicted phenotype based on the presence of AMR genes. Our results suggest that WGS data could aid to elucidate some of the expected discrepancies observed between Vitek-2 and Sensititre; for instance, the *oqxA* and *oqxB* genes conferring resistance to chloramphenicol were detected in *E. ludwigii* assemblies, and *blaSHV* gene conferring resistance to beta-lactams was detected in *K. pneumoniae* assemblies (**Table 4**). As proof of the efficacy of the approaches applied, AMR genes corresponded to resistant phenotypes to tetracyclines and chloramphenicol within total genomic as well as plasmid DNA preparations of the NR1 plasmid (control) as previously characterized (**Table 4**) [21]; however, these results varied depending on the AMR database applied as described in section 3.6 and **Table 4**.

**Table 4.** Correspondence of experimentally determined resistance phenotypes *versus* AMR genes detected in total genomic DNA^a^.

| Strain | Resistance phenotype<sup>c</sup> | Antibiotic Class | Genes detected by |  |  |
| --- | --- | --- | --- | --- | --- |
|  |  |  | ResFinder<sup>a</sup> | AMRFinder<sup>a</sup> | CARD<sup>a</sup> |
| *E. coli*<br>DU1040(NR1) | Cefalexin | Beta-lactam | ND | *blaEC* | *Amp-C* and *Amp-H Beta-lactamases* |
|  | Doxycycline | Tetracycline | *tetB, mdf(A)*<sup>d</sup> | *tetB* | *tetB* |
|  | Tetracycline | Tetracycline | *tetB, mdf(A)*<sup>d</sup> | *tetB* | *tetB, mdf(A)*<sup>d</sup> |
|  | Chloramphenicol | Phenicol | *catA1, mdf(A)*<sup>d</sup> | *catA1* | *catA1, mdf(A)* |
| *E. ludwigii*<sup>b</sup><br>LST1391B | Ampicillin | Beta-lactam | *blaACT-12* | *blaACT* | *blaACT-12* and *Amp-H Beta-lactamases* |
|  | Amoxicillin/clavulanic acid | Beta-lactam | *blaACT-12* | *blaACT* | *blaACT-12* and *Amp-H Beta-lactamases* |
|  | Cefazolin | Beta-lactam | *blaACT-12* | *blaACT* | *blaACT-12* and *Amp-H Beta-lactamases* |
|  | Cefalexin | Beta-lactam | *blaACT-12* | *blaACT-12* | *blaACT-12* and *Amp-H Beta-lactamases* |
|  | Chloramphenicol | Phenicol | ND | *oqxA, oqxB*<sup>e</sup> | ND |
| *K. pneumoniae*<sup>b</sup><br>LST1504-C2 | Ampicillin | Beta-lactam | *blaSHV-40* | *blaSHV* | *blaSHV-40* and *Amp-H Beta-lactamases* |
|  | Amoxicillin/clavulanic acid | Beta-lactam | *blaSHV-40* | *blaSHV* | *blaSHV-40* and *Amp-H Beta-lactamases* |
|  | Cefazolin | Beta-lactam | *blaSHV-40* | *blaSHV* | *blaSHV-40* and *Amp-H Beta-lactamases* |
|  | Cephalexin | Beta-lactam | *blaSHV-40* | *blaSHV* | *blaSHV-40* and *Amp-H Beta-lactamases* |

<sup>a</sup>AMR Annotation databases: AMRFinder, ResFinder and CARD. Results based on Illumina-hybrid, Nanopore-polished and Illumina-only assemblies (see Methods section and footnote of Table 1 for details of assembly pipelines).
<sup>b</sup> *Enterobacter cloacae* complex is intrinsically resistant to ampicillin, amoxicillin/clavulanic acid, and cefazolin. *K. pneumoniae* is intrinsically resistant to ampicillin.
<sup>c</sup> Susceptible phenotypes: strains were susceptible to the tested antimicrobials based on Vitek-2 and Trek Sensititre testing systems. For a complete list of susceptible and resistant phenotypes to 21 different antimicrobial agents see Table 4.
<sup>d</sup> *mdf(A)* has broad-spectrum activity but it is unclear which level of resistance it confers (Kim et al 2011).
<sup>e</sup> No resistance genes corresponding to this resistance phenotype were detected in the Nanopore-only assembly, but AMR genes were detected in Nanopore-hybrid, Illumina-only and Illumina-hybrid using the AMRFinder database.
ND: not detected

### 3.6 Comparison of AMR gene databases

To evaluate the efficiency with which AMR genes could be detected from different publicly available databases, we compared the gene symbol output obtained from each database in total genomic preparations. Most genes called by AMRFinder and ResFinder were identical (n=7), with 14 genes in total being called by AMRFinder and 9 genes in total called by ResFinder, while CARD called 88 AMR genes among the three bacterial strains (**Figure 3**; **Table S6-8**). Besides having great database resources, these results highlight the stringent search of AMRFinder and ResFinder to avoid false positives. In contrast, CARD uses less stringent cut-off thresholds increasing false positives and overcalling resistance.

**Figure 3.**
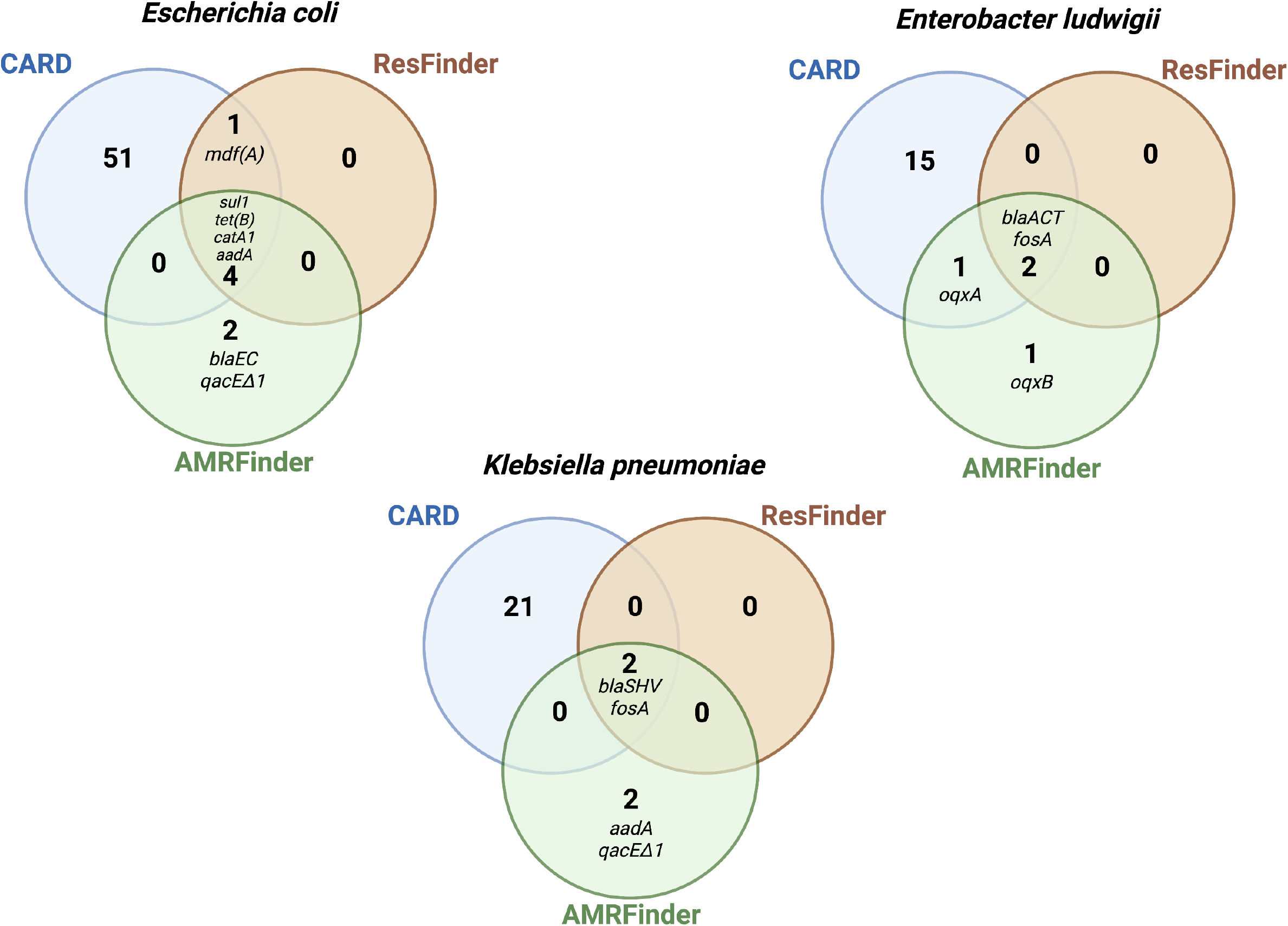
Antimicrobial resistance (AMR) genes with matches in three databases: AMRFinder, ResFinder, and Comprehensive Antibiotic Resistance Database (CARD). Venn diagrams are based on total genomic DNA sequences (Illumina-hybrid approach), and sizes do not reflect the number of AMR genes. See Table 6-8S for the description of genes listed by CARD.

We then compared AMR gene outputs of AMRFinder, ResFinder and CARD with phenotypic resistance results. AMRFinder detected *blaEC* gene in the *E. coli* strain, and *oqxA*/*oqxB* genes in *E. ludwigii*, which are consistent with the phenotypic resistance identified to beta-lactams and chloramphenicol (**Table 4, Figure 3**). In contrast, ResFinder missed these two categories of phenotypic resistance and CARD missed one category (chloramphenicol in *E. ludwigii*) (**Table 4**).

AMRFinder additionally screened for biocide and metal resistance genes identifying the *qacDeltaE1* gene (quaternary ammonium) in *K. pneumoniae* and *E. coli* (**Figure 3**), but it missed a well-characterized mercury-conferring resistance gene in the *E. coli* plasmid (control) [21].

### 3.6 Comparison of sequencing platforms and assembly approaches for detection of AMR genes

Since the accuracy of base-calling and of assembly methods influence the identification and order of the bases in the output of any sequencing platform and long- and short-read methods have technical advantages and liabilities [5] we evaluated how these different approaches affected the detection of AMR genes (**Table 5**). We observed that the Illumina-only and hybrid or Nanopore-polished approaches allowed for a more accurate detection of AMR genes in the control strain (*E. coli*) than the Nanopore-only assemblies. For instance, the *sul1* and *tet(B)* genes were not detected in Nanopore-only assemblies of *E. coli*, therefore failing to predict resistance to chloramphenicol and tetracyclines, to which *E. coli* was shown to be resistant in the phenotypic *in vitro* testing (**Tables 4 and 5**). Similarly, genes conferring resistance to chloramphenicol (*oqxA* and *oqxB*) were not detected in Nanopore-only assemblies of *E. ludwigii*, therefore, failing to predict phenotypic resistance to chloramphenicol (**Tables 4 and 5**).

**Table 5.**
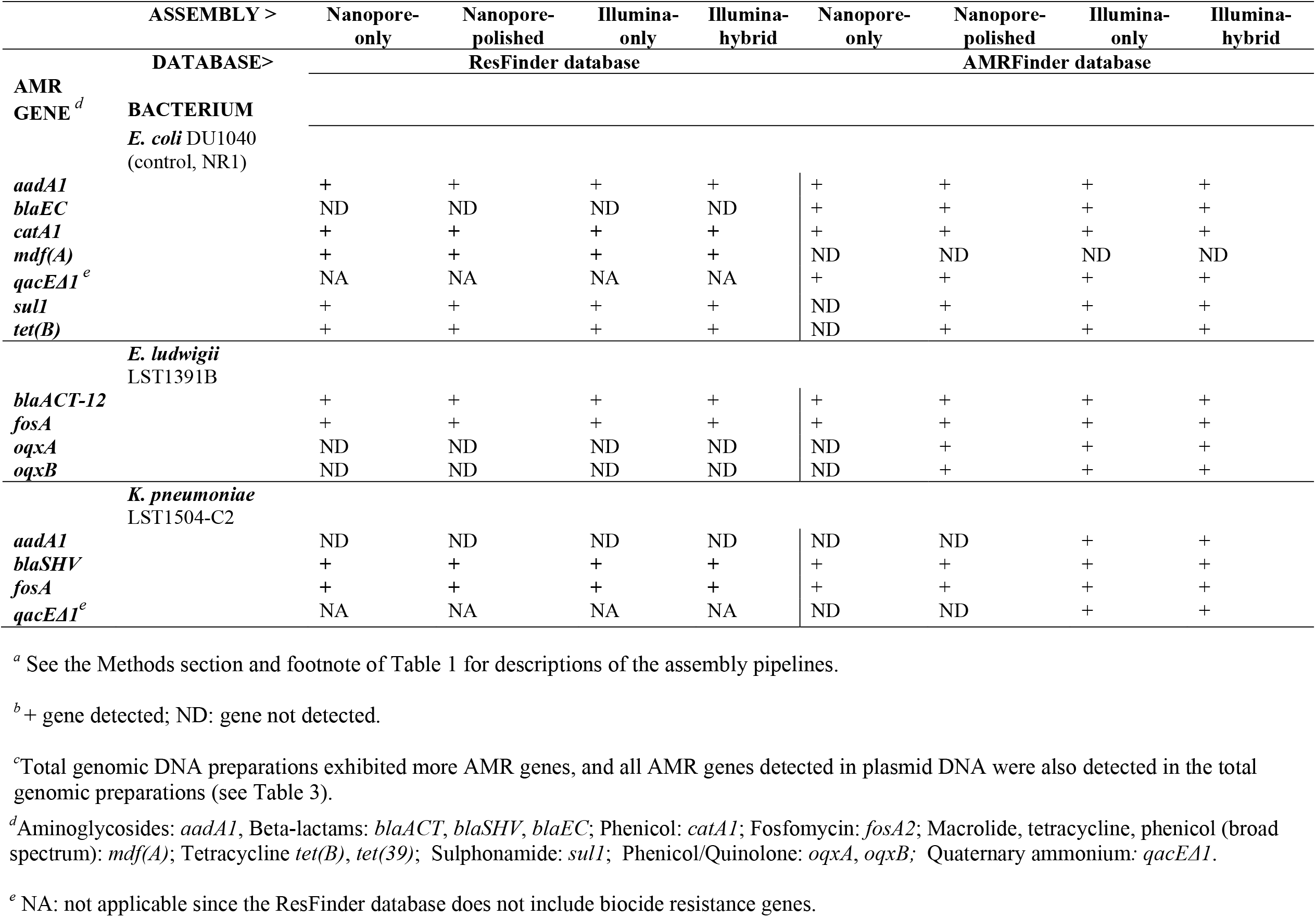
Comparison ^*a*^ of two sequencing chemistries, four assembly pipelines, and two databases for detection^*b*^ of antimicrobial resistance (AMR) genes in total genomic DNA^*c*^.

For the two test strains, *E. ludwigii* and *K. pneumoniae*, AMR genes expected to confer phenotypic resistance to chloramphenicol and beta-lactams (**Table 4**) were not detected in plasmids sequenced by Nanopore and assembled by Flye (Nanopore-only and Nanopore-polished approaches). Therefore, we further investigated if the issue was related to the type of plasmid assembler employed. We found that assembling plasmids with different long-read tools (i.e., Flye *versus* Canu) affected the detection of AMR genes (**Tables S7-8**). Genes expected to be detected in *E. ludwigii* plasmids such as *fosA* and *blaACT* were identified in assemblies from Canu but not from Flye (**Table S7**). However, in *K. pneumoniae*, either Flye or Canu failed to improve detection of a beta-lactam conferring resistance gene in plasmid-only assemblies (**Table S8**).

## 4. DISCUSSION

Whereas there has been a considerable progress in the cost and availability of WGS, integrating these technologies into routine clinical microbiology remains challenging. To our knowledge, this is a pioneer study to assess critically each step in AMR gene detection in WGS. Our results motivate more such investigations to guide identification of best practices of DNA extraction, sequencing platforms, assembly methodologies, and publicly available databases for clinically relevant bacteria, e.g. the ESKAPE list. Our results are extendable, especially since the evaluated methods are also frequently used for other bacterial pathogens.

As most of the AMR genes are in mobile elements and DNA extraction methods may impair the extraction of plasmids [11, 13], we evaluated whether a plasmid extraction step has to be incorporated in a sequencing workflow to obtain consistent identification of AMR genes. We found that a separate plasmid extraction step was not better since all AMR genes detected in the plasmid-only assemblies were also present in the total genomic DNA assemblies and corresponded in size and abundance to plasmids observed by gel electrophoresis. The literature evaluating the potential impact of DNA extraction workflows on subsequent AMR detection is limited. Salting-out kits have exhibited difficulty extracting small plasmids from *K. pneumoniae* [11] and presented impaired plasmid extraction performance in *E. coli* [13]; however, it did not influence the downstream analysis of Illumina generated WGS data [13, 37]. Here we employed a solid-phase (anion-exchange) total genomic DNA kit, Genomic-tip 20/G-Qiagen, which has been demonstrated rapidly and inexpensively to provide sufficient sequencing quality plasmid DNA [13]. Plasmid extractions can be cumbersome and not adaptable in a clinical diagnostic setting; therefore, we adapted an existing commercial extraction kit to purify plasmids directly from bacterial colonies harvested from an agar plate, resulting in a turnaround time of 6 hours, yielding a high concentration and quality of plasmid DNA meeting ONT’s DNA quality recommendations. However, despite the faster turnaround time, the extraction protocol followed by sequencing and assembling steps are still cumbersome and may not add any additional information.

We further evaluated the ability of WGS approaches to identify AMR genes associated with an experimental phenotype. Using MICs provided by automated susceptibility testing as a gold standard, we demonstrated that the presence of AMR genes was a good predictor of resistance but not a good predictor of susceptible phenotypes. Regarding resistance prediction, we showed that all categories of phenotypic resistance could be associated with AMR genes based on AMRFinder database results. Recent studies demonstrated the power to predict AMR from genomic data in clinical isolates of *E. coli, K. pneumoniae*, and *E. cloacae* complex [9, 38]. Although promising, we do not intend to suggest that WGS can replace phenotypic susceptibility testing but rather serve as a complementary method particularly useful for fastidious and slow-growing bacteria [39]. It can also be useful in cases of discrepant MIC results between different susceptibility methods as observed in *K. pneumoniae*, where the detection of *BLASHV* gene could clarify the interpretation of discrepant MICs for amoxicillin/clavulanic acid.

We observed significant variation among the three AMR databases in reporting bona fide mobile antibiotic resistances versus chromosomal housekeeping genes in which rare spontaneous resistance mutations could occur. The presence of such housekeeping genes was associated with susceptible MIC values, therefore failing to predict susceptibility. Such results may have clinical implications considering that the detection of chromosomal housekeeping genes, overcalled as “AMR genes”, could mislead clinicians to believe that the bacterium is resistant whereas it is susceptible. CARD results consisted mainly of chromosomal genes which confer transient up-regulation of efflux pumps or redox stress defense and may barely confer clinically relevant resistance. As CARD does not include mobile genes database [18], many of the transient genes from chromosomal DNA were detected instead of AMR genes from mobile elements, and many of the listed genes only have the potential to become resistant without intrinsically conferring high resistance levels. The limited stringency of CARD may impair the practicality of gene output interpretation in a clinical context. AMRFinder showed to be a valuable resource for acquired AMR genes, however, AMRFinder screening of biocide and metal resistance was not complete given that it missed the well-characterized mercury conferring resistance gene in the *E. coli* plasmid (control) [21]. Based on our data, AMRFinder and ResFinder provided easy output interpretation, low number of overcalled “AMR genes” and good prediction of resistance phenotypes. We suggest the adoption of at least two *in silico* databases in a clinical setting to ensure comparison of their outcomes to precisely identify AMR genes.

We conclude that the methods of choice may significantly influence the detection of AMR genes. The total genomic DNA preparations detected mobile element AMR genes as well as the plasmid DNA preparations, and possibly even better because many AMR genes are chromosomally borne. Nanopore-only sequences missed AMR genes and failed to predict *E. coli* and *E. ludwigii* phenotypic resistance. Illumina-only was as accurate as the hybrid assemblies but it is still slow and expensive for critical care. The pace at which microfluidics and nanochemistries are addressing the AMR detection problem may soon remedy the cost and accuracy of data acquisition challenges. For the moment, creative deployment of a mix of existing rapid data acquisition with real-time in-house data collection, processing, and modelling will improve ability of clinical laboratories to handle challenging cases rapidly and expedite patient treatment and AMR tracking.

## Data Availability

Bioinformatic pipeline shell script with dependencies, install instructions and usage instructions is available at https://github.com/iframst/HybridAMRgenotyping

Whole-genome sequences obtained from the total genomic DNA preparations on which this study is based are deposited at NCBI under the BioProject number: PRJNA624147. The plasmid sequences obtained from the plasmid DNA preparations are deposited under the BioProject number: PRJNA627408.

## Acknowledgments

We thank all the Athens Veterinary Diagnostic Laboratory technologists for exceptional technical support, especially Paula Bartlett.

## Supplementary material

**Table S1**. Concentration and purity of total genomic DNA and pure plasmid DNA preparations.

**Table S2**. Unassembled read data from Illumina Iseq and MinION Nanopore sequencing runs.

**Table S3**. Characteristics and quality assessment of four different sequencing assembly approaches by QUAST v5.0.2 (http://quast.sourceforge.net/quast).

**Table S4**. Summary of whole-genome and plasmid annotations by Prokka v.1.13.

**Table S5**. Plasmid taxonomic classification by COPLA classifier.

**Table S6**. *Escherichia coli*. Complete list of antimicrobial resistance genes detected by three different databases using two different sequencing chemistries and four different assembly approaches.

**Table S7**. *Enterobacter ludwigii*. Complete list of antimicrobial resistance genes detected by three different databases using two different sequencing chemistries and four different assembly approaches.

**Table S8**. *Klebsiella pneumoniae*. Complete list of antimicrobial resistance genes detected by three different databases using two different sequencing chemistries and four different assembly approaches.

---

Table 4 footnote reference: Kim, J.Y., Jeon, S.M., Kim, H., Park, M.S. and Kim, S.H., 2011. A contribution of MdfA to resistance to fluoroquinolones in Shigella flexneri. Osong Public Health and Research Perspectives, 2(3), pp.216–217.

